# Dynamic blue light-inducible T7 RNA polymerases (Opto-T7RNAPs) for precise spatiotemporal gene expression control

**DOI:** 10.1101/140871

**Authors:** Armin Baumschlager, Stephanie K. Aoki, Mustafa Khammash

## Abstract

Light has emerged as control input for biological systems due to its precise spatiotemporal resolution. The limited toolset for light control in bacteria motivated us to develop a light-inducible transcription system that is independent from cellular regulation through the use of an orthogonal RNA polymerase. Here, we present our engineered blue light-responsive T7 RNA polymerases (Opto-T7RNAPs) that show properties such as low leakiness of gene expression in the dark-state, high expression strength when induced with blue light, or an inducible range of more than 300-fold. Following optimization of the system to reduce expression variability, we have created a variant, which returns to the inactive dark-state within minutes, once blue light is turned off. This allows for precise dynamic control of gene expression, which is a key aspect for most applications using optogenetic regulation. The regulators were developed and tested in the bacterium *Escherichia coli*, which is a crucial cell factory for biotechnology due to its fast and inexpensive cultivation and well understood physiology and genetics. However, minor alterations should be sufficient to allow their use in other species in which the T7 RNAP polymerase and the light-inducible Vivid regulator were shown to be functional, which comprises other bacterial species and eukaryotes such as mammalian cells or yeast. We anticipate that our approach will expand the applicability of using light as an inducer for gene expression independent from cellular regulation, and allow for a more reliable dynamic control of synthetic and natural gene networks.

## INTRODUCTION

Small molecule induced gene expression systems are a key component in synthetic biology^1^ and biotechnological applications^2^. However, chemical inducers are limited in their application in space and time. Spatiotemporal control is of increasing interest, as biological systems are regulated dynamically and respond to intracellular stimuli and changes in internal states^3^. Although static perturbations, such as growth media variation and gene knockouts, have been extensively and successfully used to elucidate gene network structure and function, approaches using dynamic perturbations are providing new insight into the organizing principles of biology and the study of gene networks^3^. Dynamic regulation is also starting to be explored by metabolic engineers^4–6^. However, very recent work addresses the problem that few broadly applicable tools are available for dynamic pathway regulation, and further show that dynamic regulation can significantly increase product titers through dynamic pathway regulation^7^.

Light-based regulation is superior to conventional small-molecule inducers in this regard, displaying better temporal properties, as removal of small molecules might be challenging in scenarios such as batch or fed-batch processes. Further, it allows for spatial control of individual cells (whereas small molecules are diffusion-limited), and is minimally invasive^8^, a desired feature for basic research. These distinguishing properties of light over small molecules led to the development of numerous light-controlled devices^9–13^. Light-inducible dimerization domains were successfully exploited in two-hybrid-like systems^14^ to create optogenetic gene expression systems in eukaryotes^8–10,15^, and for reconstitution of functional proteins from their split parts^16–19^.

Dynamically light-inducible systems allow new regulation schemes, by moving the controller out of the cell and using light as input signal for control. Both biology and engineering make use of feedback control to achieve robust regulation, which in turn allows natural and engineered systems to function reliably in the face of disturbances or changing environmental conditions. However, the design of synthetic biological feedback controllers remains challenging due to the fact that biological parts do not behave as reproducibly as electronic ones. To overcome this obstacle, *in silico* feedback control was introduced by our group, which allows for electronic control of biological responses^20,21^.

Another challenge for precise control is that depending on the growth phases, nutrient conditions, and other extrinsic factors, the concentration of RNA polymerase varies, ranging from 1,800 to 10,200 molecules per cell^22^. Along with fluctuations in the ribosome concentration, this can result in changes of expression levels^23–25^ and reduce the performance of constitutive promoters. This poses a challenge to systems that require precise balances in expression levels^26^, especially when media and growth conditions change such as during industrial scale-up^27^. To decouple expression of a gene of interest from cellular RNAP concentrations, the heterologous T7 DNA-dependent RNA polymerase (T7RNAP), originating from the T7 bacteriophage^28,29^, is commonly used for protein overexpression. The polymerase shows high processivity, a high selectivity for the T7 promoter, and does not transcribe sequences from endogenous *Escherichia coli* DNA^30^. Apart from biotechnology, the enzyme has also found applications in basic biological research, such as metagenomic screening^31^. As T7RNAP-driven transcription is independent from the native *E. coli* RNAP, it allows inhibition of the native transcription machinery (e.g. with rifampicin), without affecting the orthogonal T7 transcription system, resulting in exclusive expression of T7RNAP expressed genes^30^. Further, T7RNAP variants have been engineered to recognize different promoter variants^32–35^, which allow independent expression of multiple genes. The T7RNAP has been used in yeast strains such as *Saccaromyces cerevisiae^36,37^* or *Pichia Pastoris*^38^, other bacterial species apart from *E. coli* including the nonenteric bacterium *Pseudomonas aeruginosa*^39^, and the biotechnologically relevant gram positive *Bacillus subtilis*^40^, as well as mammalian cells^41^ and higher plants^42^.

In this work, we show that the T7RNAP can be made light-inducible by splitting the polymerase into two fragments, and fusing it to photo-activatable dimerization domains. Our protein engineering strategy was guided by previous studies showing that the T7RNAP can be split^26,32,43–45^ and reconstituted through dimerization to enhance and control its function ^26^, as well as structural information about the T7RNAP^46^. We aimed to implement light-control using the heterodimeric “Magnet” domains^16^. Magnets were engineered from the small homodimerizing photoreceptor Vivid (VVD) from the filamentous fungus *Neurospora crassa*^47^ and consist of the nMag and pMag heterodimerizing protein domains, which specifically bind each other. Magnets use flavin as a chromophore for blue light-induced binding of the two domains, which is abundant in bacterial and eukaryotic cells. A similar strategy to make T7RNAP light-inducible was recently reported^48^. However, this approach uses fixed expression levels of slow-reverting photoactivatable split T7RNAP, which does not allow for easy adjustment of gene expression setpoints to experimental needs or fast reversible dynamics, a key asset of optogenetic control. Further, measurement of gene expression variability, crucial for precise cell control, was lacking as analysis was limited to bulk populations. We have addressed all these points to create a light-inducible T7RNAP expression system that allows fast and precise dynamic control. Using different protein engineering designs, our system can be brought to different setpoints of basal and maximal expression using arabinose-inducible promoters, which allows for adjustment of expression levels to experimental requirements without altering the dynamic range. This is especially important for dynamic Opto-T7RNAP versions, which have a reduced dynamic range compared to the stable (slow dark state-reverting) variants. Gene expression variability was reduced by using single cell analysis to optimize the levels of the Opto-T7RNAP domains. These improvements allowed us to develop the fast-reverting optogenetic regulator Opto-T7RNAP(563-Fast1) with precise temporal protein expression control. We further provide a set of orthogonal light-inducible T7RNAPs with different properties, that can be chosen depending on experimental needs, ranging from high expression strength for protein overexpression to high dynamic range and precise temporal control for dynamic control strategies.

## RESULTS AND DISCUSSION

### Design of light-inducible T7 RNA polymerases (Opto-T7RNAPs)

Given that the T7RNAP can be split at specific positions of its peptide chain^26,32,43–45^, and reconstitution of the functional polymerase can be enhanced through fusion of heterodimerizing coiled-coil protein structures^26^, we implemented light-control by using the light-inducible heterodimerizing domains “Magnets”^16^, due to their small size (150 amino acids) and beneficial structural features to reconstitute split-proteins. Magnets are engineered variants of the LOV photoreceptor Vivid^47^. Upon light-induction, two Vivid domains dimerize bringing the N-terminus of one domain spatially close to the C-terminus of the other binding domain (See Supplementary Figure 1). This allows for the fusion of these domains to the C-terminus of the N-terminal fragment of split proteins, and to the N-terminus of the C-terminal split fragment, reconstituting the enzyme in a spatial manner, incorporating optogenetic regulation into the T7RNAP through light-induced assembly and dissociation.

Previous studies have shown that the T7RNAP can be split at multiple positions. Therefore, our first objective was to identify which split positions allow for modifications without impairing the function of the enzyme structurally or sterically. We selected five split positions, the two previously reported positions between amino acids 179/180 and 600/601^26,32,43–45^, as well as amino acid positions 69/70, 302/303 and 563/564 (Figure 1A). We chose the latter three sites through structural analysis of a transcribing T7RNAP initiation complex^46^, which was guided by a study that identified regions^26^ at which the T7RNAP can be split. We excluded region 763–770^26^ as it contains the T7 promoter recognition loop (739–772).The chosen positions 69, 302, 563 all lie in surface-exposed flexible loops of the T7RNAP, to minimize structural and steric interference of the additional Magnet domains. Further, the amino acids at the split positions 69(A)/70(A), 302(K)/303(K), and 563(S)/564(E) are preferred residues in natural linkers^49^. We named these variants based on the amino acid they were split after to Opto-T7RNAP(69), Opto-T7RNAP(179), Opto-T7RNAP(302), Opto-T7RNAP(563) and Opto-T7RNAP(600).

**Figure 1:**
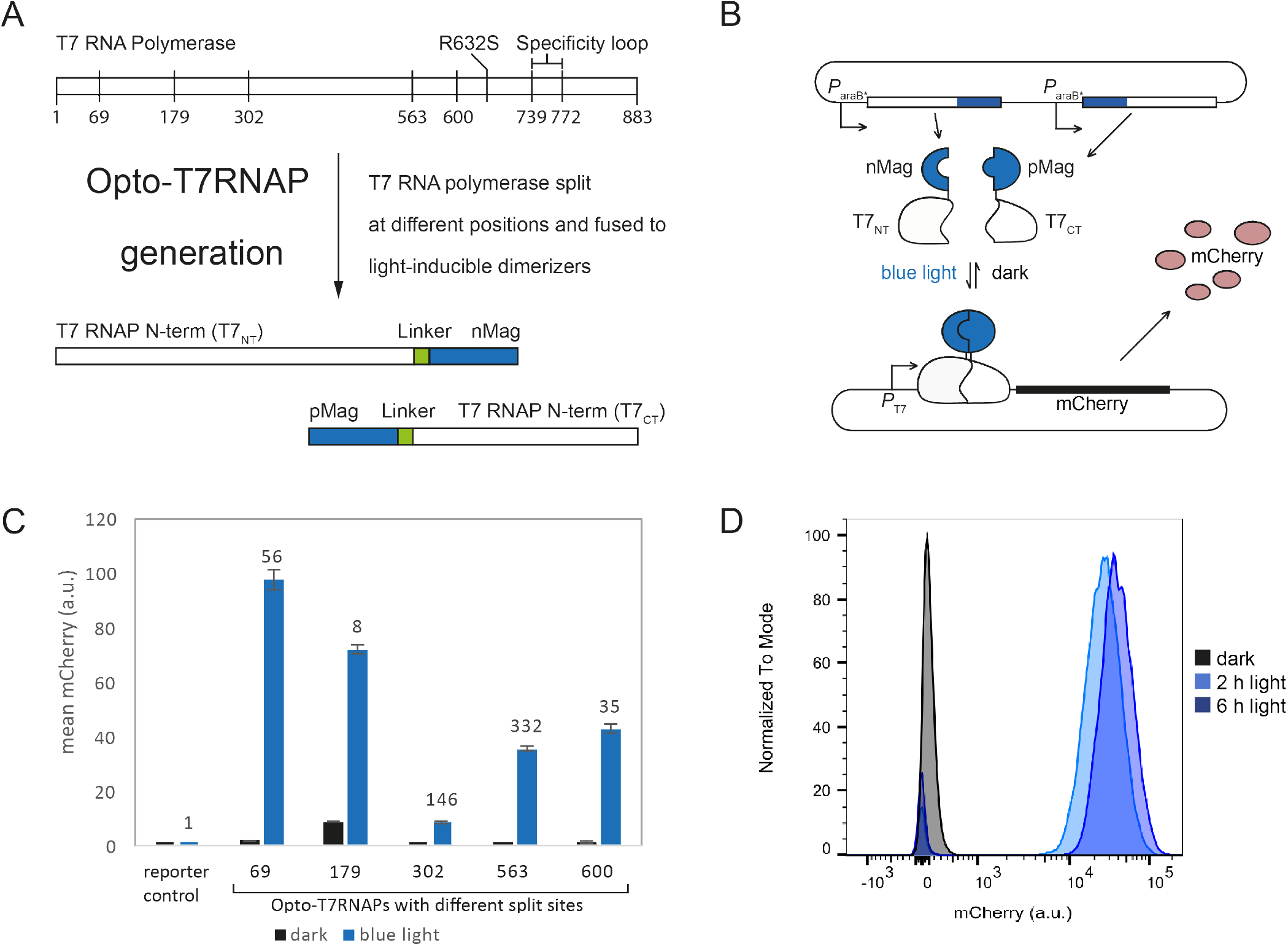
Opto-T7RNAP engineering strategy. (A) Opto-T7RNAP design. The T7RNAP is split at different positions and fused to light-inducible dimerization domains via linkers at the split site. (B) Analysis system: The Opto-T7RNAP and a reporter plasmid containing a fluorescent mCherry reporter under control of a T7 promoter is introduced into bacterial cells on two separate plasmids. Activity of Opto-T7RNAP can be measured through reporter fluorescence. (C) Steady state expression of Opto-T7RNAPs generated from the different split sites described in (A), as well as a reporter strain containing only the reporter under T7 promoter control but no T7RNAP. Raw values of mCherry a.u. and Molecules of Equivalent PE (MEPE) in Supplementary Tables 1,2 and Supplementary Figure 3. (D) Histogram of light-induced reporter gene expression in Opto-T7RNAP(563) at different time points after induction. Induction of expression is fast and unimodal. Black: uninduced, light-blue: 2h after induction, dark-blue: 6h after induction.

The N-terminal domain of the T7RNAP was fused to nMagHigh1, and the C-terminal domain of the T7RNAP fused to pMag, both using a short GGSGG-linker, schematically depicted in Figure 1A. This conformation was chosen following pretests with amino acid linkers of different length and sequence, and both conformations of nMagHigh1 and pMag fused to the N- and C-terminus of Opto-T7RNAP(179) (Data not shown). Both fusions were set under control of the *araB* promoter, excluding the CAP/CRP site, which we name *P*_araB*_. This promoter shows considerable amounts of constitutive leaky expression during log growth, and allows to increase expression though addition of the sugar arabinose (Figure 1B). To reduce expression-induced toxicity of T7RNAP, which is thought to be caused by the high processivity of the polymerase^50,51^, we introduced the mutation R632S^33^ (Figure 1A). As a reporter for Opto-T7RNAP activity, we used the red fluorescent protein mCherry, transcribed from a T7 promoter (Figure 1B). Absorption and emission of mCherry peak at 587 nm and 610 nm, which is compatible with the 460 nm activating light used for the Opto-T7RNAPs, and *vice versa*. Further, mCherry stability met the requirements of our experimental setup (Supporting Experiment “mCherry maturation assay”).

We focused on three properties that are important for expressions systems for our initial analysis of the Opto-T7RNAP variants: dark-state basal expression, light-induced expression and fold change. Dark-state basal expression provides combined information about dark-state binding of the light regulators, as well as self-assembly of the T7RNAP domains. Light-induced expression gives the maximal expression level, which is highest when the light-inducible dimerizers successfully reassemble the split T7RNAP without causing steric or structural problems. A system with a high fold change requires light-induced dimerization of the split T7RNAP for assembly to the functional enzyme, without hindering its function. We define the fold change comparing expression levels at dark and light-induced conditions, without removal of basal fluo-rescence of the bacterium or leaky expression of the reporter mCherry from the T7 promoter without T7RNAP.

We analyzed expression levels of the Opto-T7RNAPs with different split sites on the single cell level through flow cytometry of cells grown in the dark and under light induction with 329 μW/cm^2^ of 460 nm blue light (Figure 1C) during exponential growth. Opto-T7RNAP(69) reached the highest expression level, although with significant dark-state expression, resulting in a 56-fold change. Opto-T7RNAP(563) showed the highest fold change (>300-fold), with a dark-state expression of ~5-fold in the dark, and ~1’900-fold in the lit state above the reporter control. Opto-T7RNAP(302), however, showed the lowest basal expression with an induction of >140-fold in response to blue light. Gene expression is uni-modal and fast in all cases, as exemplarily shown in the histogram in Figure 1D, reaching high expression levels already after 2 h (further example Supplementary Figure 4).

It was not possible to compare the Opto-T7RNAPs to full length T7RNAP, as no cells containing both, the full length T7RNAP expressed from the *araB* promoter and mCherry under T7 promoter control, could be obtained. Protein expression can be a burden on the cell^52,53^, and mutations might arise that reduce this burden. Those variants are selected for due to reduction of cellular stress and would eventually outgrow the population due to increased fitness. The high metabolic burden and stress caused by unregulated T7-driven mCherry expression, led to a W727C mutation in the T7RNAP in all colonies after transformation into the testing strain. The T7RNAP plasmid was sequenced, confirming that it did not contain the mutation prior to transformation. This variant shows titratable T7-driven expression (Supplementary Figure 5), however does not allow comparison with the Opto-T7RNAPs.

In general, inducible gene expression systems have to fulfill different requirements, based on the applications they are used for. Heterologous protein production, for example, often requires a separation of growth and production phase. Opto-T7RNAP(69) could be used for such tasks as it allows for fast cell growth in the dark-state, and strong blue light-induced expression due to the high activity of the regulator at low regulator expression levels. If, however, the product of the T7-driven expression is toxic, a tight regulator such as Opto-T7RNAP(302) might be preferred. Opto-T7RNAP(563), provides the highest fold change. Although different variants might be relevant for different tasks, we further focused on optimizing the 69 and the 563 split Opto-T7RNAPs.

### Unequal expression of Opto-T7RNAP domains improves fold change and maximal expression level, and reduces gene expression variability

We further investigated how expression ratios of the two Opto-T7RNAP fragments influence properties important for inducible gene expression systems. Therefore, we created constructs in which we vary the RBS strength of the C-terminal Opto-T7RNAP fragment to 1.9-fold, 0.5-fold and equal predicted translation initiation rate (TIR) relative to the N-terminal Opto-T7RNAP fragment, which was kept the same (Figure 2A; TIR predictions from RBS calculator v2.0^54,55^).

**Figure 2:**
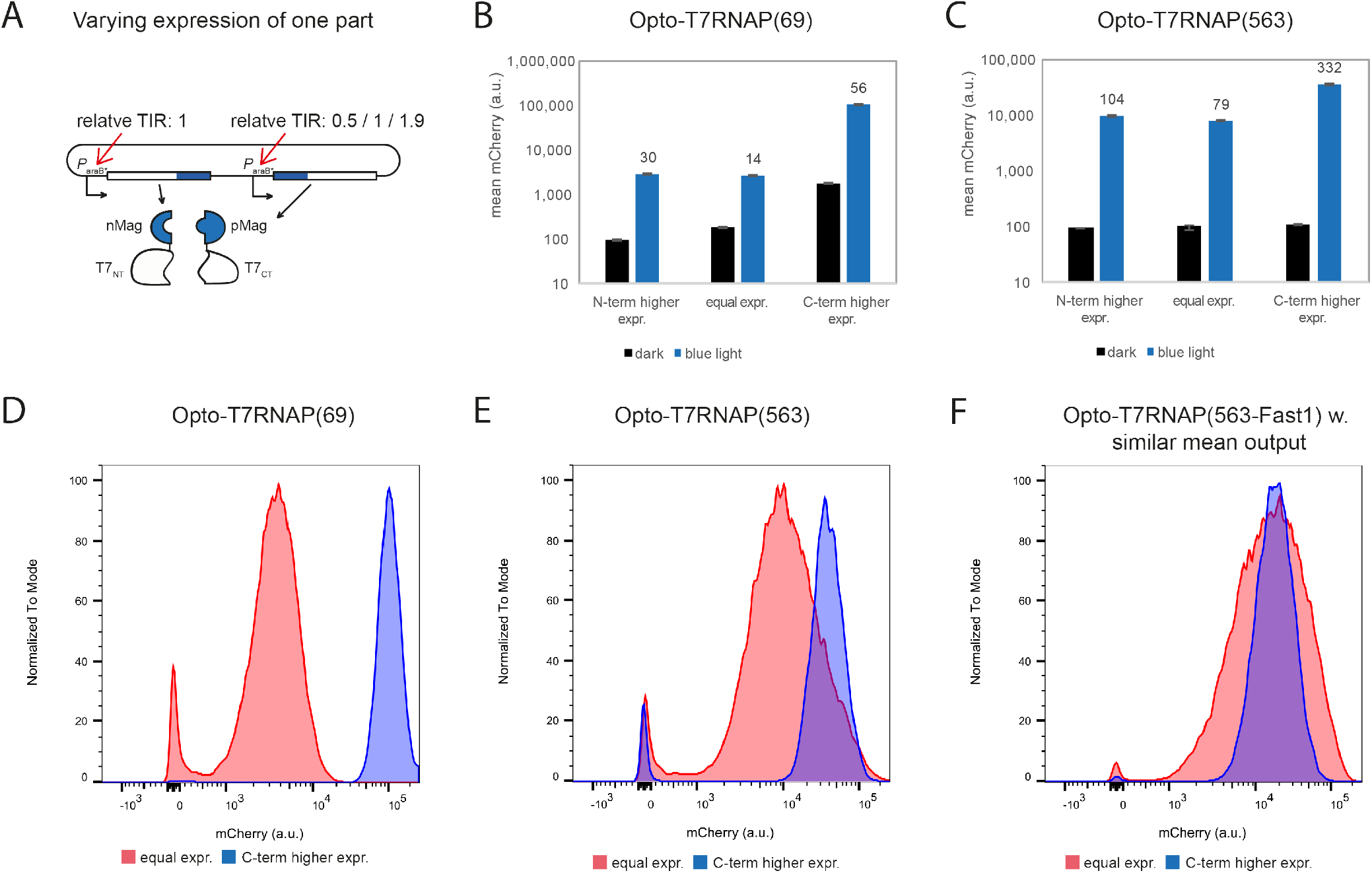
Higher expression of C-terminal Opto-T7RNAP split fragment improves fold change and expression variability. Different split fragment expression levels (A) for (B) Opto-T7RNAP(69) and (C) Opto-T7RNAP(563). Expression level of the C-terminal Opto-T7RNAP fragment (pMag fused to C-terminal T7RNAP) was set to 0.5-fold, equal, and 1.9-fold TIR of the N-terminal Opto-T7RNAP fragment (from left to right in diagrams A and B). (C) Light-induced steady state expression of equal (red) and 1.9-fold expression variants (blue) for (D) Opto-T7RNAP(69), (E) Opto-T7RNAP(563), and (F) Opto-T7RNAP(563-Fast1). Equal expression resulted in CVs of 66% for the 69 split and 137% for the 563 split variants, and were reduced through higher expression of the C-terminal fragment to CVs of 38% for the 69 split and 58% for the 563 split variants.

Expression levels above the background signal is necessary to allow for comparison of fold changes between the constructs. All Opto-T7RNAP(69) constructs showed dark state expression above background without arabinose induction, but expression levels of Opto-T7RNAP(563) with 0.5-fold and equal C-terminal fragment expression had to be increased with arabinose to generate dark state expression above background. We observed that induced expression was significantly increased when the C-terminal fragment was expressed higher. The fold change was lowest when the regulators were expressed in equal amounts, and improved with higher expression of either the N- or C-terminal fragment of both the 69 and 563 split Opto-T7RNAPs (Figure 2AB). Further, we noticed a significant decrease in mCherry expression variability, when the C-terminal fragment was over expressed (CV: 38% for Opto-T7RNAP(69), 58% for Opto-T7RNAP(563)) compared to equal expression of the domains (CV: 66% for Opto-T7RNAP(69), 137% for Opto-T7RNAP(563)) as shown in Figure 2D,E,F. Since mean expression levels can have an influence on the variability, we chose arabinose levels so that a similar light-induced mean expression level was reached for equal and higher C-terminal expressing Opto-T7RNAP(563). Also here, we observed a decrease in variability of the variant with the C-terminal fragment higher expressed (See Supplementary Figure 6). We assume that with equally expressed domains, the amount of functional Opto-T7RNAP depends on the variability of the expression of each domain. However, if the C-terminal fragment is overexpressed, the concentration of functional regulator is only dependent on the variability of the lower expressed fragment. This leads to a decreased variability of reporter gene expression.

### Dynamic light-inducible T7RNAP

One of the main assets of using optogenetics in biological systems is the ability for precise spatiotemporal control. Therefore, our aim was to develop a light-inducible expression system which reacts rapidly to changes in the light input. Since the Magnet dimerization system can only be activated, but not deactivated with light, the dynamics depend on the reversal rate to the inactive dark-state. It was reported that pMag and nMag dissociate with a half-life (*t*_1/2_) of 1.8 h^16,48^, which is not suitable for fast dynamic changes of the light input. Therefore, we used previously reported mutations in pMag that decrease dissociation time to half-lives of 4.2 min (mutation I85V for pMagFast1) and 25 s (mutations I74V and I85V for pMagFast2)^16^.

We implemented these mutations in Opto-T7RNAP(563) to create Opto-T7RNAP(563-Fast1) and Opto-T7RNAP(563-Fast2). Due to the change in regulator, we tested if unequal expression is also beneficial for these regulators, as previously described. We observed a significant reduction in the output variability similar to the slow dark-reverting regulators (Figure 2C). The fold change also increased, although less pronounced as for the stable regulators (data not shown). Therefore, we further used the conformation in which the C-terminal split fragment is expressed 1.9-fold higher than the N-terminal fragment.

To test the dynamic properties of the system, and if dark-state reversal of the regulators also leads to dissociation of the Opto-T7RNAP domains, we induced the cells with 329 μW/cm^2^ blue light for 3 h, before turning off the light, and monitored how fast reporter gene expression stops. Measurements were started during the last 45 min of light-induction, and continued for 2.5 h after light was turned off.

For the Opto-T7RNAP(563-Fast1), a slight decrease of fluorescence was observed at the first time point after 5 min, and turned into exponential decay 10-15 min after light was turned off (Figure 3A). Considering the median half-lives of mRNAs in *E. coli* of 4.7 min^56^, we observed a similar half-live to the 4.2 min that were previously reported for pMagFast1. Fluorescence of the stable variant pMag was constant for 30 min, before slowly transitioning into exponential decay (Supplementary Figure 7). Since the cells are kept in exponential growth phase, we also expect exponential decay of active Opto-T7RNAP, resulting in decrease of fluorescence. The decrease in fluorescence of the stable Opto-T7RNAP(563) after 30 min could be a combination of both dark-state reversal and dilution of active Opto-T7RNAP due to cell growth. Surprisingly, we observed a similar dynamic behavior for Opto-T7RNAP(563-Fast2) (Supplementary Figure 7), which was reported and shown to revert to the dark-state in 25 s. In the context of Opto-T7RNAPs, we did not observe an increased dark-state reversal rate of pMagFast2 compared to the stable pMag. Therefore, using pMagFast1 in Opto-T7RNAP(563-Fast1) allows for dynamic expression control.

**Figure 3:**
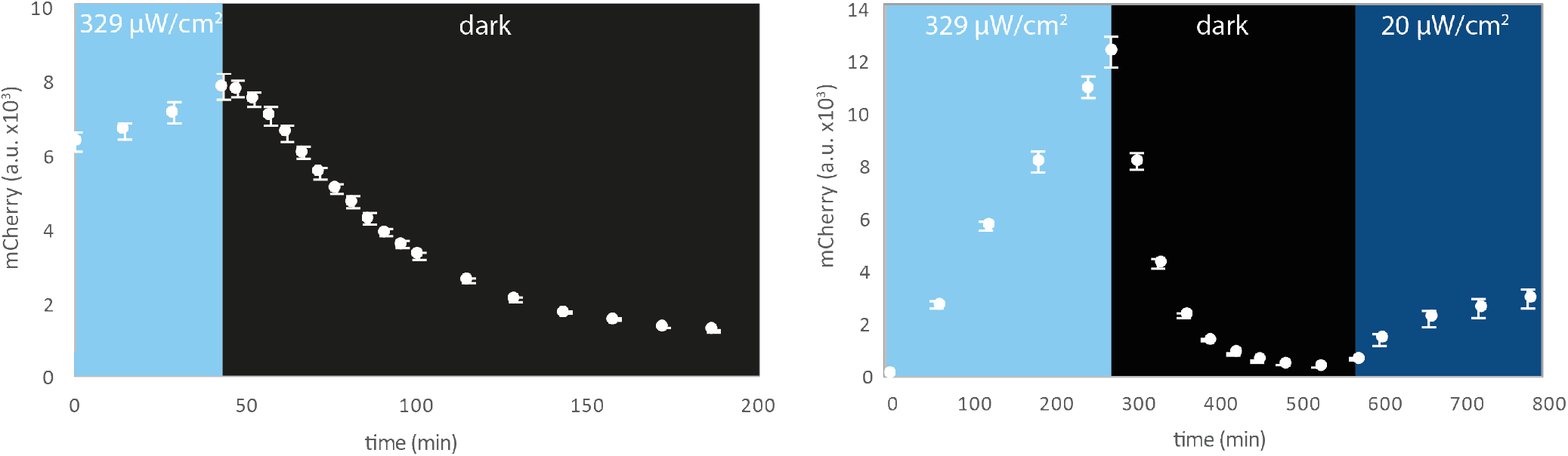
Dark-state reversal of Opto-T7RNAP(563-Fast1). Cells were induced for 3h with saturating blue light of which the last 45 min are shown, before light was turned off.

### Expression level set points and light sensitivity

Since the Opto-T7RNAP domains are under control of inducible promoters, the maximal expression of a gene of interest can be adjusted to desired values by adjusting the arabinose concentration. To allow for a titratable arabinose induction, we inserted a mutated *lacY* permease (*lacYA177C*) into the *attB* site of our testing strain BW25113^57,58^, abolishing the all-or-nothing induction of the native arabinose transporter^59^. Increased expression levels of the dynamic Opto-T7RNAP(563-Fast1), through raised arabinose concentrations, led to increased reporter expression without significant changes in the fold change (Supplementary Figure 8), as dark-state and light-induced expression increased comparably (Figure 4 left). For the stable Opto-T7RNAP(563), reporter expression was maximal with 0.1% arabinose, and decreased with 0.2% arabinose, which we suspect to be due to the additional burden of higher regulator expression and mCherry reporter expression at the maximal level of the cell (Supplementary Figure 9).

**Figure 4:**
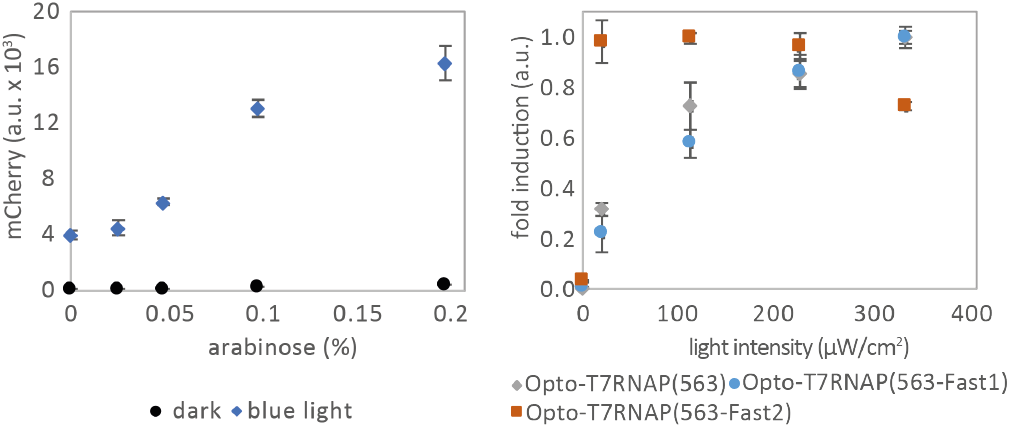
Characterization of Opto-T7RNAP expression set-points and blue light dose-response. (Left) Arabinose induction to increase Opto-T7RNAP(563-Fast1) expression levels in dark and 329 μW/cm^2^ with 460 nm light-induction conditions results in higher reporter expression with similar fold changes. (Right) Dose-response for 460 nm blue light. Fluorescence was normalized for individual constructs to allow for direct comparison.

To further characterize light sensitivity of the Opto-T7RNAP(563) variants with pMag, pMagFast1 and pMagFast2, we tested the dose-response of the three regulators to different intensities of 460 nm blue light. Opto-T7RNAP(563) and Opto-T7RNAP(563-Fast1) showed a similar dose response, with increasing expression until ~330 μW/cm^2^ (Figure 4 right). The additional I74V mutation, however, led to increased light sensitivity and maximal induction with light intensities as low as 20 μW/cm^2^, while showing low basal expression in the dark. Our experimental setup did not allow for light intensities below this value. Change in expression ratios of the Opto-T7RNAP domains, as used to optimize fold-induction (Figure 2AB), resulted in identical light sensitivities for all three regulators and does not have an influence on light sensitivity (Supplementary Figure 10).

### Fast-reverting light-inducible T7 RNA polymerase allows for fast dynamic regulation

Finally, we show that Opto-T7RNAP(563Fast1) can be used to precisely and dynamically induce gene expression. In Figure 3B we show an exemplary dynamic time course by first inducing gene expression at maximal level with 329 μW/cm^2^, resulting in high reporter expression, before shutting down the system by turning off the light at minute 270, which leads to an immediate drop of fluorescent reporter at the following time point. We then induced the system with low-intensity blue light at 20 μW/cm^2^ at minute 570, resulting in low expression of reporter protein. This demonstrates that Opto-T7RNAP(563-Fast1) allows for dynamic regulation of gene expression at different induction levels.

## CONCLUSION

Pioneering work on light-inducible gene expression was performed in *E. coli* using two-component systems^11,60,61^. In this work, we took a different approach by implementing light control into the orthogonal T7RNAP, an enzyme which is highly relevant in biotechnology and biological sciences. This was made possible by inspiring studies on the T7RNAP^26,32^. Our light-inducible system could eliminate the use of costly inducers, such as IPTG, which can be relevant for high volume microbial production systems^62,63^ and allow for precise dynamic regulation. The system could be adapted for the use in other organisms such as other bacterial species, yeast or mammalian cells, in which the T7RNAP and Magnets where shown to be functional.

With Opto-T7RNAP(563), we have created a light-inducible system that shows a >300-fold change in response to light, and fast unimodal light-induced gene expression. Further, Opto-T7RNAP(563-Fast1) allows dynamic regulation, due to fast reversion of the light-induced regulator to the dark-state when light is absent. With a fold change of >50-fold, it outperforms any previously published dynamic blue light systems (EL222: 5-fold^64^, iLID-T7: 26-fold^65^). The fast dark-state reversal also allows using pulse width modulation (PWM), which was previously applied to optogenetically-controlled systems^66^, and might be useful for additional fine-tuning of expression levels.

Optimized expression levels of the two Opto-T7RNAP domains allow for a more precise regulation due to reduced gene expression variability. Further, the system can be brought to desired set points of minimal and maximal expression in both the dynamic and the stable system. This allows for adaptation of the system to experimental requirements.

The presented Opto-T7RNAPs all respond to blue light. However, due to the modularity of the system, the blue light sensitive Magnets could be exchanged with regulators that dimerize in response to other wavelengths. As T7RNAP variants have been engineered to recognize different promoter sequences^32–35^, Opto-T7RNAPs that respond to different wavelengths could be used in the same cell for multi-chromatic control of multiple genes. Considering regulation schemes, which use multiple optogenetic regulators for differential gene expression, regulators that activate at a certain wavelength and revert quickly in the dark, such as the Magnet system used here, might be preferable to systems which need light of a different wavelength for dark-state reversion as they leave a greater portion of the light spectrum available for other optogenetic regulators.

The absorbance spectrum of Vivid^8,47^ suggests that light with wavelengths >500 nm should not activate the regulator, which would make it possible to combine this system with available light-inducible systems of different wavelength, such as the green/red controllable CcaS-CcaR, which is activated with 535 nm, and inactivated with 670 nm, or the red/far-red controlled Cph8-OmpR, which is activated with 703 nm or in the dark and inactivated with 650 nm^60^.

We further found that Opto-T7RNAP(563-Fast2) shows a high sensitivity to blue light, which might be preferable, if only low light levels can be provided for activation.

The Opto-T7RNAPs will also serve as an excellent screening system to engineer the properties of the light-inducible dimerizing Magnets towards lowering dark-state interaction and increasing light-induced binding affinity and dark-state reversal rates through rational protein design and directed evolution, as it provides an easy screen for transcriptional output.

We further showed, that unequal expression of the two Opto-T7RNAP domains can lead to reduced gene expression variability, a feature which is desirable for precise control, as well as higher gene expression productivity, as the fraction of cells in a high expression state can be increased. It has to be elucidated if this finding also holds for other light-inducible split proteins.

## MATERIALS AND METHODS

### Strains and Media

*Escherichia coli* DH5αZ1^67^ was used for all cloning. For characterization we used *E. coli* strain AB360, that we derived from *E. coli* BW25113^57,58^, which we modified by integrating *lacYA177C* into the *attB* site^8^. This transforms the all-or-nothing induction of the native arabinose transporter towards titratable arabinose induction^59^. Relevant for this study is that the strains contain the transcription factor *araC*, while arabinose metabolizing genes *araBAD* are deleted. Autoclaved LB-Miller medium was used for strain propagation. Sterile-filtered M9 medium supplemented with 0.2% casamino acids, 0.4% glucose and 0.001% thiamine was used for all gene expression experiments. Antibiotics were used if necessary for plasmid maintenance in concentrations of 100 μg/ml ampicillin, 25 μg/ml chloramphenicol, and 50 μg/ml kanamycin.

### Plasmids and genetic parts

Plasmids were transformed using a one-step preparation protocol of competent *E. coli* ^68^. We used the pZ-series modular vectors^67^ containing the pSC101 ori with *repA,* and *cat* for chloramphenicol resistance, as basis for plasmids containing the Opto-T7RNAPs. The vector was modified for expression of two proteins, so that it contained two arabinose promoters *P*_araB_* (*araB* promoter from Guzman et al.^69^, excluding CAP binding site), followed by the Opto-T7RNAP fusions and *rrnB* T1 terminator sequences (BBa_B0010), which are each flanked by unique cut sites to allow for modular exchange of promoters and RBSs. The N-terminal Opto-T7RNAP fusions can be exchanged between BglII and AvrII, its promoters with AatII and BglII, the C-terminal Opto-T7RNAP can be exchanged between either NsiI (for pAB164, pAB166, pAB170) or PacI (for all other plasmids) and XbaI, and it promoters between AscI and either NsiI or PacI. (See plasmid map in Supplementary Figure 11) The empty reporter control strain contains the same plasmid, with *araC* instead of the Opto-T7RNAP. The T7 polymerase split fragments were amplified from pTARA^70^ (Addgene plasmid # 31491). The plasmid contained a missense mutation at residue position 823 which was corrected to the original sequence with OEPCR. We further introduced the mutation R632S^33^ to all T7RNAP constructs, also through OEPCR and primers with sequence AGT for amino acid position 632. For the T7 promoter driven mCherry reporter plasmid, we used pETM6-mCherry^71^ (a gift from Mattheos Koffas) and inserted the T7 promoter with the same sequence also used in pTHSSD_8^26^ with a strong RBS on synthesized oligos between AvrII and NdeI. The TIRs for expression level studies of the Opto-T7RNAP fragments were calculated using the RBS calculator v2.0^54,55^, with the resulting mRNA structures and TIRs shown in Supplementary Figure 12. All resulting strains are listed in Supplementary Table 3 and genetic parts are listed in Supplementary Table 4 which were cloned using standard molecular biology protocols^72,73^.

### Integration of *lacYA177C* into *E. coli* BW25113

We used λ integrase expressed from pJW27 to integrate *lacYA177C* into the *attB* site of BW25113^57,58^, using plasmid pSKA27 (unpublished, sequence available upon request) containing *lacYA177C*, FRT-flanked *kanR* from pKD13 ligated into XbaI cut pFL503^59^ and sequence identical to genome regions for *attB* integration. pSKA27 was cut with NotI and the 4229 bp band gel purified and circularized before transformation into pJW27 containing cells. For integration, pJW27 was transformed into *E. coli* BW25113^57,58^, and selected at 30°C on LB-Agar plates containing chloramphenicol, for expression of λ integrase. A single colony was used to inoculate 5 ml LB broth containing chloramphenicol, and grown at 30°C in a water bath with shaking. The cells were then moved to 42°C for 15 min, before incubating on ice for 15 min. Cells were transformed with the integration construct using the previously described transformation protocol. The kanamycin resistance cassette can be removed by expressing FLP-recombinase following the protocol of Datsenko and Wanner^58^ by transforming pCP20, and select for chloramphenicol and ampicillin at 30°C. The kanamycin resistance was kept in all strains for this study.

### T7RNAP-induced gene expression during steady state growth

All experiments were performed in biological triplicates for data shown in the main text, as well as data in the supplementary material unless explicitly noted. Expression of the Opto-T7RNAPs was chosen, so that dark-state expression levels of all tested constructs were above the reporter control (cells containing just the T7 promoter controlled mCherry without T7RNAP), which allows for comparison of fold change between the constructs.

All experiments consist of time course induction experiments performed with cells in exponential growth phase to allow for reliable comparison of expression levels. The single cell flow cytometry data shown are steady state, or close to steady state expression levels of cells in exponential growth phase. The corresponding time courses are provided in the Supplementary Figures 13–17. Further, we provide molecules of equivalent fluorophore for the expression levels of the different Opto-T7RNAP variants in Supplementary Figure 3, which allows researchers to compare our mCherry expression levels, independent from the technical equipment.

Overnight cultures inoculated from glycerol stocks, originating from single colonies, were grown in M9 medium containing chloramphenicol, ampicillin and kanamycin at 37°C with 200 rpm shaking in lightproof black tubes (Greiner bio-one). The samples were diluted 1:25000 in 7 ml fresh M9 medium in sterile polystyrol tubes (Sarstedt, ethanol washed, and capped with aluminum foil). The tubes were wrapped in aluminum foil to prevent outside light from other experiments reaching the cells, leaving only the bottom uncovered for induction with LED light. All experiments were performed in a dark room.

For expression experiments, cells were incubated at 37°C in a water bath (Julabo, ED (v.2) THERM60) and custom made LED lights similar as described by Milias-Argeitis^21^, with the difference that led lights were placed under the tubes in the water bath, and LEDs with 460 nm wavelength (40 lm, 700 mA; LED Engin Inc.) were used. Tubes were placed on a multi position magnetic stirrer (Thermo scientific, Telesystem 15), for stirring with teflon covered 3 x 8 mm magnetic stirrers (Huberlab AG) at 900 rpm (Supplementary Figure 18).

Samples were grown without light induction for at least 2.5 h before the experiment was started to allow adaptation towards log growth phase in fresh M9 media, containing arabinose for some experiments, and steady state expression of the regulators. Samples were taken every 60 min for steady state expression experiments or at shorter intervals for dynamic experiments. Cells were kept below an OD_600_ of 0.03 and therefore in logarithmic growth phase through manual dilution with fresh medium of the respective arabinose concentration.

### mCherry maturation assay

We determined maturation of the fluorescent protein mCherry for our experimental setting to 90 min for full maturation with translation and transcription inhibition (Supporting Experiment “Transcription and translation inhibition”, Supplementary Figure 19). Sample was added to the inhibition solution in equal volumes, resulting in a final inhibitor concentration of 250 μg/ml rifampicin (Sigma-Aldrich Chemie GmbH) and 25 μg/mL tetracycline (Sigma-Aldrich Chemie GmbH). The transcription and translation inhibition solution contained 500 μg/mL rifampicin, 50 μg/mL tetracycline in phosphate buffered saline (Sigma-Aldrich Chemie GmbH, Dulbecco’s Phosphate Buffered Saline) and filtered using a 0.2 μm syringe filter (Sartorius). The inhibition solution was precooled on ice. After sample was added, the solution was incubated on ice for at least 30 min, to allow for diffusion of the antibiotics into the cell and inhibition of transcription and translation in aluminum foil-covered 96-well U-bottom plates (Thermo Scientific Nunc). In between wells of the 96-well plate that contained sample, we added 200 μL PBS to include an additional wash step, which reduces carryover of sample to undetectable levels using the BD^TM^ High Throughput Sampler of the flow cytometer. After incubation on ice for 30 min, the sample were transferred to a 37°C incubator for 90 min for mCherry maturation. Then the cells were put on ice until measurement through flow cytometry.

### Flow cytometry measurement

Cell fluorescence was characterized on a BD LSR Fortessa containing a 561nm laser with BD™ High Throughput Sampler attached, and analyzed using FlowJo vX (TreeStar). mCherry fluorescence was characterized with a 561nm (100 mW) laser and 610/20 nm band pass, 600 nm long pass filter. At least 10,000 events were recorded in a two-dimensional forward and side scatter gate, which was drawn by eye and corresponding to the experimentally determined size of the testing strain at logarithmic growth (Supplementary Figure 20). Gating for analysis was also determined by eye, and was kept constant for analysis of all experiments and used for calculations of the median and CV using the same software.

## ASSOCIATED CONTENT

### Supporting Information

Supporting experiments and figures (PDF)

## AUTHOR INFORMATION

### Author Contributions

AB conceived the project, and MK supervised the project. AB designed and performed the experiments and analyzed the data. SKA cloned the *lacYA177C* integration plasmid pSKA27 and provided experimental advice. AB, SKA, and MK wrote the manuscript. All authors have given approval to the final version of the manuscript.

### Notes

The authors declare no competing financial interest.

## ACKNOWLEDGMENT

Marc Rullan and Peter Buchmann for help with the light setup. Dr. Gupta Ankit, Dr. Corentin Briat, Dr. Andreas Milias-Argeitis and Dirk Benzinger for helpful discussions. AB is part of the Life Science Zurich graduate school.

## ABBREVIATIONS

RNAP, DNA-dependent RNA polymerase; RBS, ribosome binding site; TIR, translation initiation rate; a.u. arbitrary units

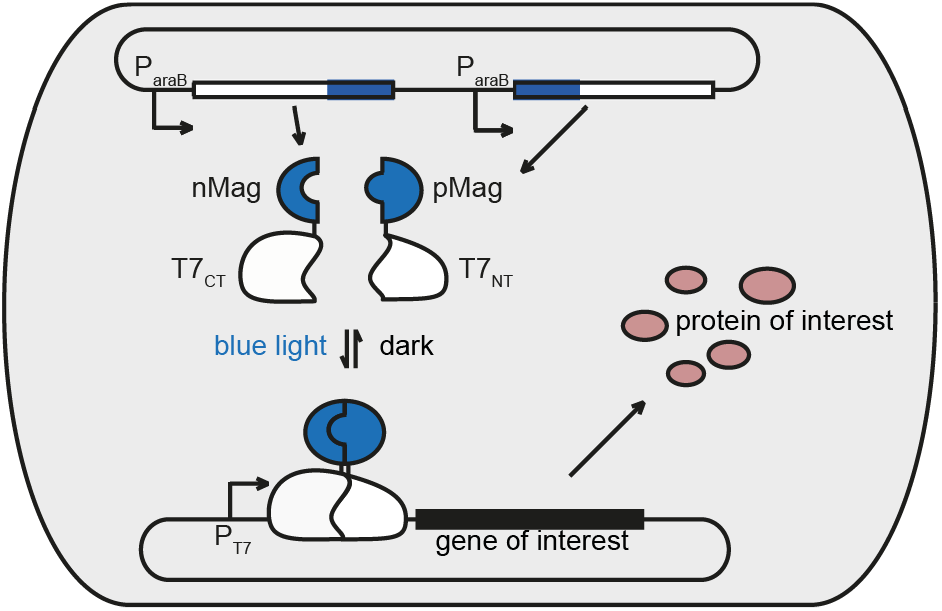

## REFERENCES

(1) Khalil, A. S. S., and Collins, J. J. J. (2010) Synthetic biology: applications come of age. Nat. Rev. Genet. 11, 367–379.

(2) Brautaset, T., Lale, R., and Valla, S. (2009) Positively regulated bacterial expression systems. Microb. Biotechnol. 2, 15–30.

(3) Castillo-Hair, S. M., Igoshin, O. a, and Tabor, J. J. (2015) How to train your microbe: methods for dynamically characterizing gene networks. Curr. Opin. Microbiol. 24, 113–123.

(4) Brockman, I. M., and Prather, K. L. J. (2015) Dynamic metabolic engineering: New strategies for developing responsive cell factories. Biotechnol. J. 10, 1360–1369.

(5) Torella, J. P., Ford, T. J., Kim, S. N., Chen, A. M., Way, J. C., and Silver, P. a. (2013) Tailored fatty acid synthesis via dynamic control of fatty acid elongation. Proc. Natl. Acad. Sci. U. S. A. 110, 11290–5.

(6) Holtz, W. J., and Keasling, J. D. (2010) Engineering Static and Dynamic Control of Synthetic Pathways. Cell 140, 19–23.

(7) Gupta, A., Reizman, I. M. B., Reisch, C. R., and Prather, K. L. J. (2017) Dynamic regulation of metabolic flux in engineered bacteria using a pathway-independent quorum-sensing circuit. Nat. Biotechnol. 3.

(8) Müller, K., Naumann, S., Weber, W., and Zurbriggen, M. D. (2015) Optogenetics for gene expression in mammalian cells. Biol. Chem. 396, 145–152.

(9) Müller, K., Engesser, R., Schulz, S., Steinberg, T., Tomakidi, P., Weber, C. C., Ulm, R., Timmer, J., Zurbriggen, M. D., and Weber, W. (2013) Multi-chromatic control of mammalian gene expression and signaling. Nucleic Acids Res. 41.

(10) Pathak, G. P., Strickland, D., Vrana, J. D., and Tucker, C. L. (2014) Benchmarking of Optical Dimerizer Systems. ACS Synth. Biol. 3, 832–838.

(11) Levskaya, A., Chevalier, A. A., Tabor, J. J., Simpson, Z. B., Lavery, L. A., Levy, M., Davidson, E. A., Scouras, A., Ellington††, A. D., Marcotte, E. M., and Voigt, C. A. (2005) Synthetic biology: engineering Escherichia coli to see light. Nature 438, 441–442.

(12) Shimizu-Sato, S., Huq, E., Tepperman, J. M., and Quail, P. H. (2002) A light-switchable gene promoter system. Nat. Biotechnol. 20, 1041–4.

(13) Williams, S. C. P., and Deisseroth, K. (2013) Optogenetics. Proc. Natl. Acad. Sci. 110, 16287–16287.

(14) Fields, S., and Song, O. (1989) A novel genetic system to detect protein-protein interactions. Nature 340, 245–246.

(15) Niu, J., Ben Johny, M., Dick, I. E., and Inoue, T. (2016) Following Optogenetic Dimerizers and Quantitative Prospects. Biophys. J. 1–9.

(16) Kawano, F., Suzuki, H., Furuya, A., and Sato, M. (2015) Engineered pairs of distinct photoswitches for optogenetic control of cellular proteins. Nat. Commun. 6, 6256.

(17) Nihongaki, Y., Kawano, F., Nakajima, T., and Sato, M. (2015) Photoactivatable CRISPR-Cas9 for optogenetic genome editing. Nat. Biotechnol. 33.

(18) Taslimi, A., Zoltowski, B., Miranda, J. G., Pathak, G. P., Hughes, R. M., and Tucker, C. L. (2016) Optimized second-generation CRY2–CIB dimerizers and photoactivatable Cre recombinase. Nat. Chem. Biol. 1–8.

(19) Kawano, F., Okazaki, R., Yazawa, M., and Sato, M. (2016) A photoactivatable Cre-loxP recombination system for optogenetic genome engineering. Nat. Chem. Biol. 12, 1059–1064.

(20) Milias-Argeitis, A., Summers, S., Stewart-Ornstein, J., Zuleta, I., Pincus, D., El-Samad, H., Khammash, M., and Lygeros, J. (2011) In silico feedback for in vivo regulation of a gene expression circuit. Nat. Biotechnol. 29, 1114–1116.

(21) Milias-Argeitis, A., Rullan, M., Aoki, S. K., Buchmann, P., and Khammash, M. (2016) Automated optogenetic feedback control for precise and robust regulation of gene expression and cell growth. Nat. Commun. 7, 12546.

(22) Bremer, H., and Dennis, P. P. (1987) Modulation of Chemical Composition and Other Parameters of the Cell by Growth Rate. Escherichia coli Salmonella Cell. Mol. Biol. 2, 1527–1542.

(23) Keren, L., Zackay, O., Lotan-Pompan, M., Barenholz, U., Dekel, E., Sasson, V., Aidelberg, G., Bren, A., Zeevi, D., Weinberger, A., Alon, U., Milo, R., and Segal, E. (2013) Promoters maintain their relative activity levels under different growth conditions. Mol. Syst. Biol. 9, 701.

(24) de Vos, D., Bruggeman, F. J., Westerhoff, H. V., and Bakker, B. M. (2011) How molecular competition influences fluxes in gene expression networks. PLoS One 6.

(25) Klumpp, S., Zhang, Z., and Hwa, T. (2009) Growth Rate-Dependent Global Effects on Gene Expression in Bacteria. Cell 139, 1366–1375.

(26) Segall-Shapiro, T. H., Meyer, A. J., Ellington, A. D., Sontag, E. D., and Voigt, C. a. (2014) A “resource allocator” for transcription based on a highly fragmented T7 RNA polymerase. Mol. Syst. Biol. 10, 742.

(27) Moser, F., Broers, N. J., Hartmans, S., Tamsir, A., Kerkman, R., Roubos, J. A., Bovenberg, R., and Voigt, C. A. (2012) Genetic circuit performance under conditions relevant for industrial bioreactors. ACS Synth. Biol. 1, 555–564.

(28) Studier, F. W. (1991) Use of bacteriophage T7 lysozyme to improve an inducible T7 expression system. J. Mol. Biol. 219, 37–4.

(29) Chamberlin, M., McGrath, J., and Waskell, L. (1970) New RNA polymerase from Escherichia coli infected with bacteriophage T7. Nature 228, 227–231.

(30) Tabor, S. (2001) Expression Using the T7 RNA Polymerase/Promoter System. Curr. Protoc. Mol. Biol. Chapter 16, Unit16.2.

(31) Lussier, F. X., Chambenoit, O., Côté, A., Hupé, J. F., Denis, F., Juteau, P., Beaudet, R., and Shareck, F. (2011) Construction and functional screening of a metagenomic library using a T7 RNA polymerase-based expression cosmid vector. J. Ind. Microbiol. Biotechnol. 38, 1321–1328.

(32) Shis, D. L., and Bennett, M. R. (2013) Library of synthetic transcriptional AND gates built with split T7 RNA polymerase mutants. Proc. Natl. Acad. Sci. U. S. A. 110, 5028–33.

(33) Temme, K., Hill, R., Segall-Shapiro, T. H., Moser, F., and Voigt, C. A. (2012) Modular control of multiple pathways using engineered orthogonal T7 polymerases. Nucleic Acids Res. 40, 8773–8781.

(34) Carlson, J. C., Badran, A. H., Guggiana-Nilo, D. A., and Liu, D. R. (2014) Negative selection and stringency modulation in phage-assisted continuous evolution. Nat Chem Biol 10, 216–222.

(35) Ellefson, J. W., Meyer, A. J., Hughes, R. a, Cannon, J. R., Brodbelt, J. S., and Ellington, A. D. (2014) Directed evolution of genetic parts and circuits by compartmentalized partnered replication. Nat. Biotechnol. 32, 97–101.

(36) Benton, B. M., Eng, W. K., Dunn, J. J., Studier, F. W., Sternglanz, R., and Fisher, P. A. (1990) Signal-mediated import of bacteriophage T7 RNA polymerase into the Saccharomyces cerevisiae nucleus and specific transcription of target genes. Mol. Cell. Biol. 10, 353–360.

(37) Chen, W., Tabor, S., and Struhl, K. (1987) Distinguishing between mechanisms of eukaryotic transcriptional activation with bacteriophage T7 RNA polymerase. Cell 50, 1047–1055.

(38) Hobl, B., Hock, B., Schneck, S., Fischer, R., and Mack, M. (2013) Bacteriophage T7 RNA polymerase-based expression in Pichia pastoris. Protein Expr. Purif. 92, 100–104.

(39) Brunschwig, E., and Darzins, A. (1992) A two-component T7 system for the overexpression of genes in Pseudomonas aeruginosa. Gene 111, 35–41.

(40) Conrad, B., Savchenko, R. S., Breves, R., and Hofemeister, J. (1996) A T7 promoter-specific, inducible protein expression system for Bacillus subtilis. Mol. Gen. Genet. 250, 230–236.

(41) Elroy-Stein, O., and Moss, B. (1990) Cytoplasmic expression system based on constitutive synthesis of bacteriophage T7 RNA polymerase in mammalian cells. Proc. Natl. Acad. Sci. U. S. A. 87, 6743–6747.

(42) McBride, K. E., Schaaf, D. J., Daley, M., and Stalker, D. M. (1994) Controlled expression of plastid transgenes in plants based on a nuclear DNA-encoded and plastid-targeted T7 RNA polymerase. Proc. Natl. Acad. Sci. U. S. A. 91, 7301–7305.

(43) Ikeda, R. A., and Richardson, C. C. (1987) Interactions of a proteolytically nicked RNA polymerase of bacteriophage T7 with its promoter. J. Biol. Chem. 262, 3800–8.

(44) Ikeda, R. A., and Richardson, C. C. (1987) Enzymatic properties of a proteolytically nicked RNA polymerase of bacteriophage T7. J. Biol. Chem. 262, 3790–9.

(45) Muller, D. K., Martin, C. T., and Coleman Joseph E. (1988) Processivity of Proteolytically Modified Forms of T7 RNA Polymerase. Biochemistry 27, 5763–5771.

(46) Cheetham, G. M., and Steitz, T. a. (1999) Structure of a transcribing T7 RNA polymerase initiation complex. Science 286, 2305–2309.

(47) Zoltowski, B. D., Schwerdtfeger, C., Widom, J., Loros, J. J., Bilwes, A. M., Dunlap, J. C., Crane, B. R., and The. (2007) Conformational Switching in the Fungal Light Sensor Vivid. Science (80-.). 316, 1054–1057.

(48) Han, T., Chen, Q., and Liu, H. (2016) Engineered photoactivatable genetic switches based on the bacterium phage T7 RNA polymerase. ACS Synth. Biol. acssynbio.6b00248.

(49) Chen, X., Zaro, J. L., and Shen, W. C. (2013) Fusion protein linkers: Property, design and functionality. Adv. Drug Deliv. Rev. 65, 1357–1369.

(50) Iost, I., Guillerez, J., and Dreyfus, M. (1992) Bacteriophage T7 RNA polymerase travels far ahead of ribosomes in vivo. J. Bacteriol. 174, 619–622.

(51) Miroux, B., and Walker, J. E. (1996) Over-production of proteins in Escherichia coli: mutant hosts that allow synthesis of some membrane proteins and globular proteins at high levels. J. Mol. Biol. 260, 289–298.

(52) Malakar, P., and Venkatesh, K. V. (2012) Effect of substrate and IPTG concentrations on the burden to growth of Escherichia coli on glycerol due to the expression of Lac proteins. Appl. Microbiol. Biotechnol. 93, 2543–2549.

(53) Dekel, E., and Alon, U. (2005) Optimality and evolutionary tuning of the expression level of a protein. Nature 436, 588–592.

(54) Espah Borujeni, A., Channarasappa, A. S., and Salis, H. M. (2014) Translation rate is controlled by coupled trade-offs between site accessibility, selective RNA unfolding and sliding at upstream standby sites. Nucleic Acids Res. 42, 2646–2659.

(55) Salis, H. M., Mirsky, E. A., and Voigt, C. A. (2009) Automated design of synthetic ribosome binding sites to control protein expression. Nat. Biotechnol. 27, 946–50.

(56) Bernstein, J. a, Khodursky, A. B., Lin, P.-H., Lin-Chao, S., and Cohen, S. N. (2002) Global analysis of mRNA decay and abundance in Escherichia coli at single-gene resolution using two-color fluorescent DNA microarrays. Proc. Natl. Acad. Sci. U. S. A. 99, 9697–702.

(57) Baba, T., Ara, T., Hasegawa, M., Takai, Y., Okumura, Y., Baba, M., Datsenko, K. A., Tomita, M., Wanner, B. L., and Mori, H. (2006) Construction of Escherichia coli K-12 in-frame, single-gene knockout mutants: the Keio collection. Mol. Syst. Biol. 2, 2006.0008.

(58) Datsenko, K. A., and Wanner, B. L. (2000) One-step inactivation of chromosomal genes in Escherichia coli K-12 using PCR products. Proc. Natl. Acad. Sci. U. S. A. 97, 6640–5.

(59) Bowers, L. M., Lapoint, K., Anthony, L., Pluciennik, A., and Filutowicz, M. (2004) Bacterial expression system with tightly regulated gene expression and plasmid copy number. Gene 340, 11–18.

(60) Schmidl, S. R., Sheth, R. U., Wu, A., and Tabor, J. (2014) Refactoring and Optimization of Light-Switchable Escherichia coli Two-Component Systems.

(61) Tabor, J. J., Levskaya, A., and Voigt, C. a. (2011) Multichromatic control of gene expression in escherichia coli. J. Mol. Biol. 405, 315–324.

(62) Lo, T.-M., Chng, S. H., Teo, W. S., Cho, H.-S., and Chang, M. W. (2016) A Two-Layer Gene Circuit for Decoupling Cell Growth from Metabolite Production. Cell Syst. 3, 133–143.

(63) Briand, L., Marcion, G., Kriznik, A., Heydel, J. M., Artur, Y., Garrido, C., Seigneuric, R., and Neiers, F. (2016) A self-inducible heterologous protein expression system in Escherichia coli. Sci. Rep. 6, 33037.

(64) Jayaraman, P., Devarajan, K., Chua, T. K., Zhang, H., Gunawan, E., and Poh, C. L. (2016) Blue light-mediated transcriptional activation and repression of gene expression in bacteria. Nucleic Acids Res. gkw548.

(65) Pu, J., Boltz, J. Z.-, and Dickinson, B. C. (2017) Evolution of a split RNA polymerase as a versatile biosensor platform. Nat. Chem. Biol. 1–27.

(66) Davidson, E. A., Basu, A. S., and Bayer, T. S. (2013) Programming microbes using pulse width modulation of optical signals. J. Mol. Biol. 425, 4161–4166.

(67) Lutz, R., and Bujard, H. (1997) Independent and tight regulation of transcriptional units in escherichia coli via the LacR/O, the TetR/O and AraC/I1-I2 regulatory elements. Nucleic Acids Res. 25, 1203–1210.

(68) Chung, C. T., Niemela, S. L., and Miller, R. H. (1989) One-step preparation of competent Escherichia coli: Transformation and storage of bacterial cells in the same solution (recombinant DNA). Pnas 86, 2172–2175.

(69) Guzman, L. L. M., Belin, D., Carson, M. J., Beckwith, J., Luz-Maria Guzman Michael J. Carson, and Jon Beckwith, D. B., and Luz-Maria Guzman Michael J. Carson, and Jon Beckwith, D. B. (1995) Tight Regulation, Modulation, and High-Level Expression by Vectors Containing the Arabinose PBAD Promoter. J. Bacteriol. 177, 4121–4130.

(70) Wycuff, D. R., and Matthews, K. S. (2000) Generation of an AraC-araBAD Promoter-Regulated T7 Expression System. Anal. Biochem. 277, 67–73.

(71) Jones, J. A., Vernacchio, V. R., Lachance, D. M., Lebovich, M., Fu, L., Shirke, A. N., Schultz, V. L., Cress, B., Linhardt, R. J., and Koffas, M. A. G. (2015) ePathOptimize: A Combinatorial Approach for Transcriptional Balancing of Metabolic Pathways. Sci. Rep. 5, 11301.

(72) Sambrook, J. (2001) Molecular cloning: a laboratory manual 3.ed. Cold Spring Harbor Laboratory Press, Cold Spring Harbor, NY.

(73) Gibson, D. G., Young, L., Chuang, R.-Y., Venter, J. C., Hutchison, C. A., and Smith, H. O. (2009) Enzymatic assembly of DNA molecules up to several hundred kilobases. Nat. Methods 6, 343–5.

